# *In vitro* and *In vivo* characterization of NOSO-502, a novel inhibitor of bacterial translation

**DOI:** 10.1101/329946

**Authors:** Emilie Racine, Patrice Nordmann, Lucile Pantel, Matthieu Sarciaux, Marine Serri, Jessica Houard, Philippe Villain-Guillot, Anthony Demord, Carina Vingsbo Lundberg, Maxime Gualtieri

## Abstract

Antibacterial activity screening of a collection of *Xenorhabdus* strains led to the discovery of the Odilorhabdins, a novel antibiotic class with broad-spectrum activity against Gram-positive and Gram-negative pathogens. Odilorhabdins inhibit bacterial translation by a novel mechanism of action on ribosomes. A lead-optimization program identified NOSO-502 as a promising candidate. NOSO-502 has MIC values ranging from 0.5 to 4 μg/ml against standard *Enterobacteriaceae* strains and carbapenem-resistant *Enterobacteriaceae* (CRE) isolates that produce KPC, AmpC, or OXA enzymes and metallo-β-lactamases. In addition, this compound overcomes multiple chromosome-encoded or plasmid-mediated resistance mechanisms of acquired resistance to colistin. It is effective in mouse systemic infection models against *E. coli* EN122 (ESBL) or *E. coli* ATCC BAA-2469 (NDM-1), achieving an ED_50_ of 3.5 mg/kg and 1-, 2- and 3-log reductions in blood burden at 2.6, 3.8, and 5.9 mg/kg, respectively, in the first model and 100% survival in the second, starting with a dose as low as 4 mg/kg. In a UTI model of *E. coli* UTI89, urine, bladder and kidney burdens were reduced by 2.39, 1.96, and 1.36 log_10_ CFU/ml, respectively, after injecting 24 mg/kg. There was no cytotoxicity against HepG2, HK-2, or HRPT cells, no inhibition of hERG-CHO or Nav 1.5 -HEK current, and no increase of micronuclei at 512 μM. NOSO-502, a compound with a novel mechanism of action, is active against *Enterobacteriaceae*, including all classes of CRE, has a low potential for resistance development, shows efficacy in several mouse models, and has a favorable *in vitro* safety profile.

## INTRODUCTION

Antibiotic-resistant infections are spreading around the world. The urgent need to discover new families of antibacterial agents to counter the threat of drug-resistant infection is widely recognized. The U.S. Centers for Disease Control and Prevention (CDC) recently published a report outlining the top 18 drug-resistant threats. Two were classified as “urgent” in terms of threat level: carbapenem-resistant *Enterobacteriacea*e (CRE) and *Clostridium difficile* (1). Carbapenems are broad-spectrum β-lactam antibiotics saved for the treatment of the most serious infections. CRE have become resistant to all or nearly all antibiotics available and cause many types of serious infection, such as those of the respiratory tract, urinary tract, abdomen and bacteremia (2). The CDC estimates that 9,300 healthcare-associated infections are caused each year in the United States by the two most common types of CRE, carbapenem-resistant *Klebsiella* species and *Escherichia coli* species, causing approximately 600 deaths (1). In China, among the 664 CRE cases reported in 2015 in 25 hospitals, most were caused by *K. pneumoniae* (73.3%), *E. coli* (16.6%), or *E. cloacae* (7.1%) and the overall mortality rate was 33.5% (2).

Antibacterial activity screening of a collection of *Xenorhabdus* strains led to the discovery of the Odilorhabdins, a novel antibiotic class with broad-spectrum activity against Gram-positive and Gram-negative pathogens (3). Odilorhabdins inhibit bacterial translation by a novel mechanism of action on ribosomes (3). Their chemical tractability made them suitable for a lead optimization program by medicinal chemistry that led to the preclinical candidate NOSO-502 (Fig. 1).

**Figure 1.**
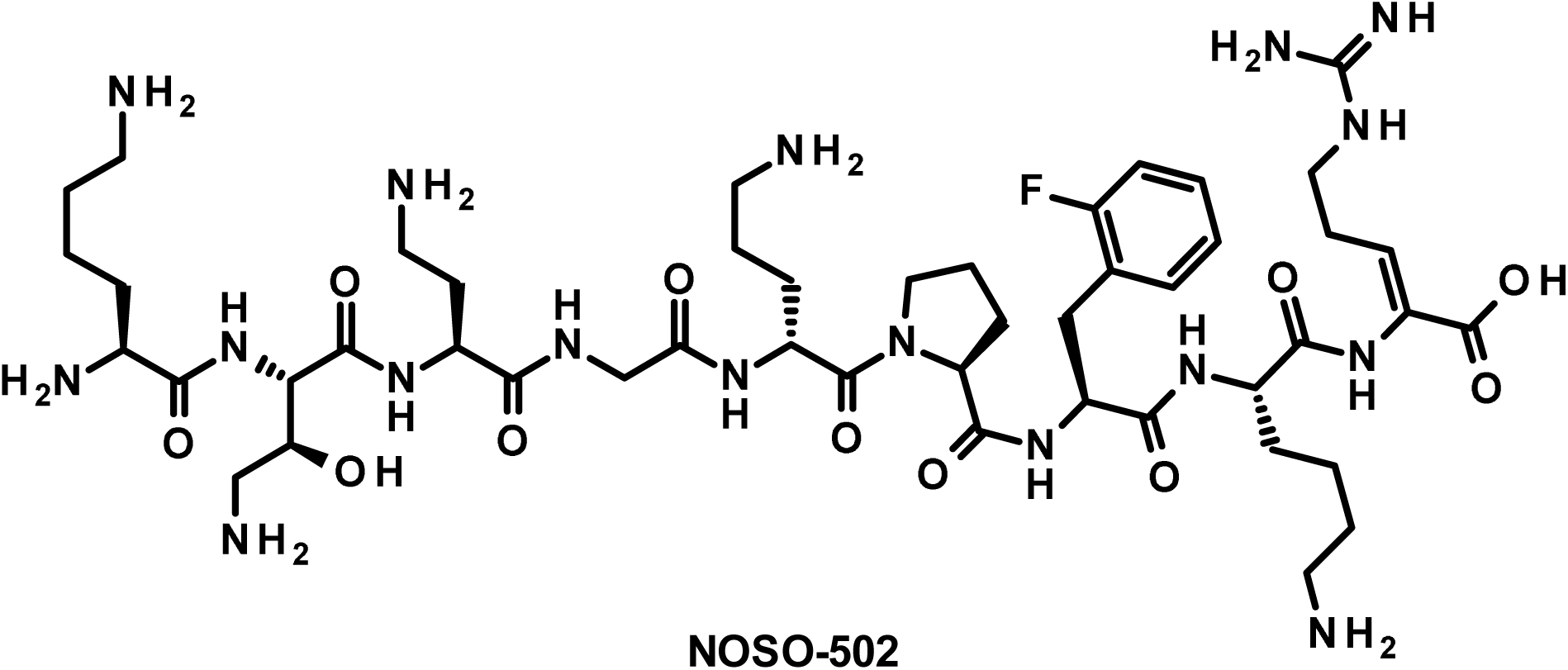
Chemical structure of NOSO-502.

We report the *in vitro* and *in vivo* characterization of NOSO-502. The data demonstrate that NOSO-502 is active against a panel of Gram-positive and Gram-negative bacteria, including carbapenem-resistant and polymyxin-resistant strains, and exhibits promising *in vivo* activity in various murine infection models, a favorable *in vitro* safety profile, and a low potential for resistance development.

## RESULTS

### NOSO-502 exhibits potent antibacterial activity

The antibacterial activity spectrum of NOSO-502 was assessed by testing a panel of Gram-positive and Gram-negative wild-type strains. The compound was active against Gram-negative pathogens of the *Enterobacteriaceae* family, such as *E. coli* or *K. pneumoniae*, with MIC values between 0.5 and 4 μg/ml, as well as *S. maltophilia*. In comparison, the MIC values of NOSO-502 against *P. aeruginosa* and *A. baumannii* were > 64 μg/ml. For Gram-positive species, NOSO-502 was more active against *Staphylococci* than *Enterococcus* or *Streptococcus* strains (Table 1).

**Table 1.** Bacterial susceptibility profile of NOSO-502 against reference bacterial strains. NOS: NOSO-502, CIP: ciprofloxacin, GEN: gentamicin, PMB: polymyxin B, IPM: imipenem, TGC: tigecycline.

| Strain | MIC (µg/ml) of antibiotics |  |  |  |  |  |
| --- | --- | --- | --- | --- | --- | --- |
|  | NOS | CIP | GEN | IPM | TGC | PMB |
| *Citrobacter freundii* DSM 30039 | 2 | ≤0.125 | 0.5 | 1 | 1 | 0.5 |
| *Citrobacter kozeri* DSM 4595 | 2 | ≤0.125 | 0.25 | 4 | 1 | 0.25 |
| *Enterobacter aerogenes* DSM 30053 | 2 | ≤0.125 | 0.25 | 2 | 1 | 0.5 |
| *Enterobacter cloacae* DSM 14563 | 2 | ≤0.125 | 0.5 | 1 | 4 | 1 |
| *Escherichia coli* ATCC 25922 | 4 | ≤0.125 | 1 | 0.25 | 0.25 | 0.5 |
| *Klebsiella pneumoniae* ATCC 43816 | 1 | ≤0.125 | 0.25 | 1 | 2 | 1 |
| *Serratia marcescens* DSM 17174 | 4 | ≤0.125 | 0.5 | 2 | 4 | >32 |
| *Acinetobacter baumannii* ATCC 19606 | >64 | 2 | 16 | 0.5 | 1 | 0.5 |
| *Pseudomonas aeruginosa* DSM 1117 | >64 | 1 | 1 | 2 | >8 | 1 |
| *Stenotrophomas maltophilia* ATCC 13637 | 16 | 1 | 4 | >64 | 0.5 | 1 |
| *Enterococcus faecalis* DSM 2570 | >64 | 2 | 16 | 1 | 0.25 | >32 |
| *Enterococcus faecium* DSM 20477 | 64 | 16 | 8 | 4 | 0.125 | >32 |
| *Staphylococcus aureus* ATCC 29213 | 1 | 0.25 | 0.5 | ≤0.125 | 0.5 | 16 |
| *Staphylococcus epidermidis* ATCC 12228 | 0.25 | 0.25 | ≤0.125 | ≤0.125 | 0.5 | 16 |
| *Streptococcus pneumoniae* DSM 2134 | 64 | 1 | 8 | ≤0.125 | 0.125 | >32 |

The compound was also tested against a recent panel of *Enterobacteriaceae* clinical isolates. MIC_90_ values were between 2 and 8 μg/ml against *E. coli, K. pneumoniae, Enterobacter cloacae*, and *Citrobacter freundii*. The antibacterial activity of NOSO-502 was conserved against fluoroquinolone-, aminoglycoside-, and polymyxin B-resistant strains of the panel (Table 2).

**Table 2.** MIC_90_ of NOSO-502 and comparators against a panel of recent clinical bacterial strains. NOS: NOSO-502, CIP: ciprofloxacin, GEN: gentamicin, PMB: polymyxin B. MIC_50_ and MIC_90_ were calculated for populations >10 isolates.

| Organsim (No. of isolates) | Antibiotic | Range | MIC (µg/ml) |  |
| --- | --- | --- | --- | --- |
|  |  |  | 50% | 90% |
| *Citrobacter freundii* (16) | NOS | 1-4 | 2 | 2 |
|  | CIP | 0.008->1 | 0.03 | >1 |
|  | GEN | 0.5->32 | 1 | >32 |
|  | PMB | 0.25-1 | 0.5 | 1 |
| *Enterobacter cloacae* (13) | NOS | 1-4 | 1 | 2 |
|  | CIP | 0.016->1 | >1 | >1 |
|  | GEN | 1->32 | 32 | >32 |
|  | PMB | 0.5-16 | 0.5 | 8 |
| *Escherichia coli* (101) | NOS | 2-32 | 4 | 8 |
|  | CIP | 0.008->1 | 0.03 | >1 |
|  | GEN | 0.5->32 | 1 | 2 |
|  | PMB | 0.25-32 | 0.5 | 1 |
| Ciprofloxacin-resistant *E. coli* (19) | NOS | 2-32 | 4 | 8 |
|  | CIP | >1 | >1 | >1 |
|  | GEN | 0.13->32 | 0.5 | >32 |
|  | PMB | 0.5-32 | 0.5 | 1 |
| Gentamicin-resistant *E. coli* (6) | NOS | 4 | - | - |
|  | CIP | 0.25-1 | - | - |
|  | GEN | 32->32 | - | - |
|  | PMB | 0.25-32 | - | - |
| Polymyxin B-resistant *E. coli* (2) | NOS | 4-8 | - | - |
|  | CIP | 1->1 | - | - |
|  | GEN | 1->32 | - | - |
|  | PMB | 4-32 | - | - |
| *Klebsiella pneumoniae* (56) | NOS | 0.5-16 | 1 | 2 |
|  | CIP | 0.008->1 | 0.5 | >1 |
|  | GEN | 0.5->32 | 1 | >32 |
|  | PMB | 0.25-32 | 0.5 | 4 |
| Ciprofloxacin-resistant *K. pneumoniae* (27) | NOS | 0.5-16 | 1 | 2 |
|  | CIP | >1 | >1 | >1 |
|  | GEN | 0.13->32 | 32 | >32 |
|  | PMB | 0.5->32 | 0.5 | 1 |
| Gentamicin-resistant *K. pneumoniae* (16) | NOS | 0.5-2 | 1 | 2 |
|  | CIP | 0.5->1 | >1 | >1 |
|  | GEN | 32->32 | >32 | >32 |
|  | PMB | 0.5-1 | 0.5 | 1 |

MIC values of NOSO-502 were determined against selected CRE and colistin-resistant isolates. The CRE strains tested produce KPC enzymes (Ambler class A carbapenemase-producing strains), metallo-β-lactamases, such as NDM, VIM, or IMP (Ambler class B carbapenemase-producing strains), AmpC (Ambler class C carbapenem-resistant strains), and OXA-48 enzymes (Ambler class D carbapenemase-producing strains). NOSO-502 exhibited potent activity against all carbapenemase-producing *Enterobacteriaceae* strains (Table 3) and overcame multiple mechanisms of colistin acquired resistance (chromosome-encoded mutations or deletions of *pmrA, pmrB, mgrB,* or *phoQ* genes or expression of *mcr-1, mcr-2*, or *mcr-3* genes), except mechanisms involving mutations of the *crrB* gene (Table 4).

**Table 3.** Activity of NOSO-502 and comparators against carbapenem-resistant *Enterobacteriaceae* strains. NOS: NOSO-502, CIP: ciprofloxacin, GEN: gentamicin, IPM: imipenem, TGC: tigecycline, PMB: polymyxin B.

|  |  | MIC (μg/ml) of antibiotics |  |  |  |  |  |
| --- | --- | --- | --- | --- | --- | --- | --- |
| Strain | β-lactamase content | NOS | CIP | GEN | IPM | TGC | PMB |
| Ambler class A carbapenemase |  |  |  |  |  |  |  |
| *Escherichia coli* PSP | KPC-2 + TEM-1 + OXA-1 | 2 | 16 | >64 | 8 | 0.5 | 0.5 |
| *Escherichia coli* COL | KPC-2 + TEM-1 + CTX-M9 | 2 | >64 | >64 | 8 | 0.5 | 0.5 |
| *Escherichia coli* MIN | KPC-3 + OXA-9 | 4 | <0.25 | 0.5 | 8 | 0.25 | 0.5 |
| *Klebsiella pneumoniae* ATCC BAA-1905 | KPC-2 | 1 | >64 | 32 | >64 | 4 | 0.5 |
| *Klebsiella pneumoniae* A33504 | KPC-2+ SHV-11 + TEM-1 + CTX-M-2 + OXA-9 | 1 | 32 | >64 | 16 | 2 | 0.5 |
| *Klebsiella pneumoniae* ATCC BAA-1904 | KPC-3 | 2 | 0.25 | 16 | 32 | 2 | 0.5 |
| *Enterobacter cloacae* KBM15 | KPC-2 + TEM-1 + OXA-9 | 1 | 32 | 8 | 32 | 8 | 0.5 |
| Ambler class B carbapenemase |  |  |  |  |  |  |  |
| *Escherichia coli* BAA-2469 | NDM-1 | 2 | 16 | >64 | 32 | 1 | 0.25 |
| *Escherichia coli* BAA-2471 | NDM-1 | 4 | >64 | >64 | >64 | 1 | 0.25 |
| *Escherichia coli* MON | NDM-5 + CTX-M15 + TEM-1 | 4 | >64 | 0.5 | 32 | 0.5 | 0.5 |
| *Escherichia coli* GAL | NDM-6 + OXA-1 + CTX-M15 | 2 | >64 | 2 | 32 | 0.5 | 0.25 |
| *Escherichia coli* EGB957 | VIM-1 + OXA-48 + TEM-1 + CMY-4 + OXA-1 | 4 | >64 | >64 | >64 | 1 | 0.5 |
| *Klebsiella pneumoniae* ATCC BAA-2146 | NDM-1 + CTX-M15 + TEM-1 + CMY-6 + OXA-1 + SHV-1 | 0.5 | >64 | >64 | >64 | 32 | 1 |
| *Klebsiella pneumoniae* NCTC 13443 | NDM-1 | 1 | >64 | >64 | >64 | 4 | 0.5 |
| *Klebsiella pneumoniae* LAM | NDM-4 + SHV-11 + CTX-M15 | 1 | >64 | >64 | 32 | 2 | 0.5 |
| *Klebsiella pneumoniae* NCTC 13439 | VIM-1 | 1 | 16 | 1 | 32 | 2 | 0.5 |
| *Enterobacter cloacae* 3047 | NDM-1 + CTX-M15 + TEM-1 + OXA-1 | 1 | 2 | 32 | 16 | 4 | 0.5 |
| Ambler class C carbapenem-resistant |  |  |  |  |  |  |  |
| *Enterobacter cloacae* 10.72 | AmpC overexpressed + TEM-1 + OXA-1 | 1 | >64 | >64 | 4 | 8 | 0.5 |
| *Citrobacter freundii* MAU | AmpC overexpressed + TEM-3 | 1 | >64 | 2 | 4 | 8 | 0.5 |
| Ambler class D carbapenemase |  |  |  |  |  |  |  |
| *Escherichia coli* DOV | OXA-48 + TEM-1 + CTX-M15 + OXA-1 | 2 | >64 | 32 | 4 | 2 | 0.5 |
| *Klebsiella pneumoniae* NCTC 13442 | OXA-48 | 1 | 4 | 0.25 | 16 | 2 | 0.5 |
| *Klebsiella pneumoniae* DUB | OXA-48 + CTX-M15 + TEM-1 + SHV-1 + OXA-1 + CMY-2 | 0.5 | >64 | >64 | 16 | 4 | 0.5 |
| *Enterobacter cloacae* YAM | OXA-48 + CTX-M15 + TEM-1 + OXA-1 + DHA-1 | 2 | >64 | >64 | 4 | 4 | 1 |
| *Enterobacter cloacae* BEU | OXA-48 + CTX-M15 + SHV-12 + TEM-1 + OXA-1 + DHA-1 | 1 | 32 | >64 | 16 | 16 | 16 |

**Table 4.**
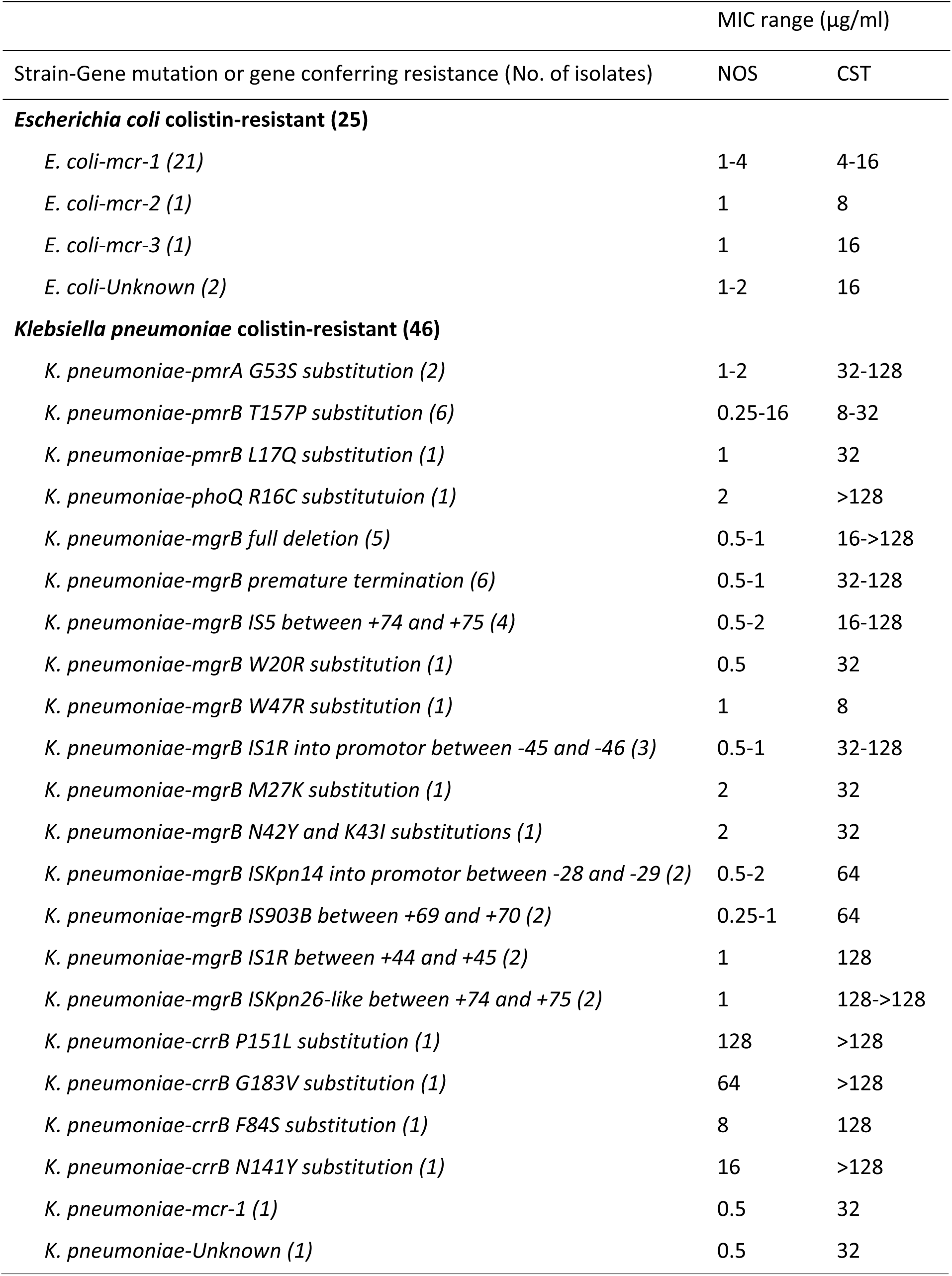
Bacterial susceptibility profile of NOSO-502 against colistin-resistant strains. NOS: NOSO-502, CST: colistin.

NOSO-502 had rapid bactericidal activity against *E. coli* ATCC 25922 and *K. pneumoniae* ATCC 43816, causing a 3-log decrease in CFU/ml at 1 h (4× and 8× MIC) (Fig. 2). We observed regrowth of *E. coli* at 4× MIC. Such regrowth at 24 h is not uncommon and has previously been reported for bactericidal antimicrobials, such as ciprofloxacin against *E. coli* (4).

**Figure 2.**
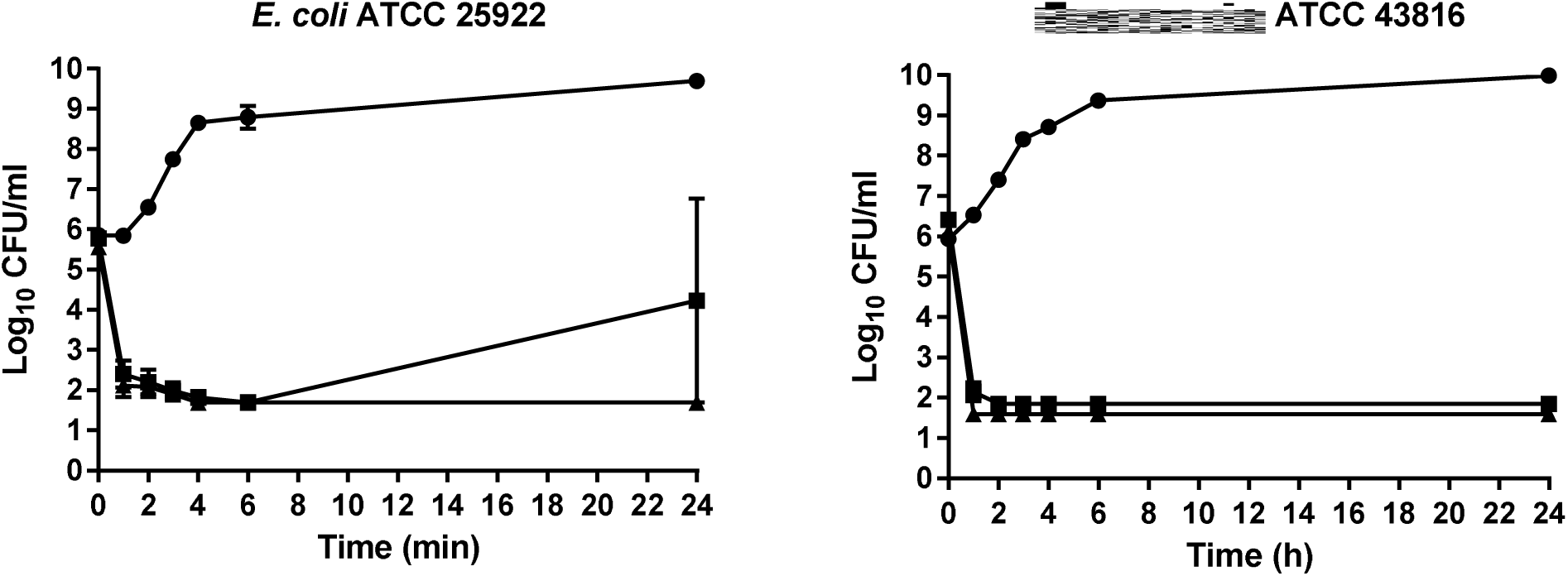
Bactericidal activity of NOSO-502 at 4× and 8× MIC against *E. coli* ATCC 25922 and *K. pneumoniae* ATCC 43816. Closed circles, drug-free control; closed squares, NOSO-502 at 4× MIC; closed triangles, NOSO-502 at 8× MIC. Experiments were performed in triplicate. Each symbol represents the mean and error bars indicate the standard error of the mean.

The propensity of bacteria to develop resistance to NOSO-502 was assessed by determining the spontaneous frequency of resistance (FoR) to the compound with *E. coli* ATCC 25922 and *K. pneumoniae* ATCC 43816. Mutants of *E. coli* resistant to 4× MIC (16 μg/ml) or 8× MIC (32 μg/ml) of NOSO-502 were isolated at a frequency of 3.0× 10^-9^ and <5.0× 10^-10^, respectively. The frequency of resistance of *K. pneumoniae* was 2.4× 10^-9^ at 4× MIC (4 μg/ml) and < 7× 10^-10^ at 8× MIC (8 μg/ml).

### NOSO-502 has a good *in vitro* safety profile

The potential nephrotoxicity of NOSO-502 was assessed in cells derived directly from human kidney tissue, human renal proximal tubular epithelial cells (HRPTEpiC), and HK-2. A multiplexed assay with HRPTEpiC cells was used to assess cellular stress *in vitro* induced by NOSO-502. Three parameters were measured: a decrease in cell viability, the expression of heat shock protein 27 (HSP27), and the level of kidney injury molecule-1 (KIM-1). A decrease in cell viability is a very sensitive marker to detect general toxicity but is not sufficient to predict nephrotoxicity, whereas increases in the level of biomarkers, such as KIM-1 or HSP27, are well correlated with dose levels of known nephrotoxic compounds (5, 6). HSP27 is expressed in response to cellular stress to block the apoptotic pathway. KIM-1 is a well-accepted marker of renal proximal tubule injury. NOSO-502 showed no cytotoxicity to HRPT cells and the molecule did not significantly increase (five-fold) KIM-1 or HSP27 levels at concentrations up to 100 μM. Polymyxin B and gentamicin, used as comparators in this study, showed different toxicity profiles. Polymyxin B was cytotoxic at low concentrations (IC_50_ = 11.8 μM) and induced a significant increase of KIM-1 and HSP27 levels at 12.1 and 9.7 μM, respectively. Gentamicin was not cytotoxic and did not increase KIM-1 levels at concentrations up to 100 μM but induced a five-fold increase of HSP27 levels at 22.4 μM. NOSO-502 did not show any effect on HK-2 cell viability at concentrations up to 512 μM (0% inhibition from 16 to 256 μM and 9.4% inhibition at 512 μM).

The cardiotoxic effect of NOSO-502 was evaluated using the automated patch clamp human ether-a-go-go related gene (hERG) potassium channel assay. This test is now accepted as an early predictor of potential cardiotoxicity and is used routinely at an early stage in the drug discovery process. NOSO-502 did not significantly inhibit hERG currents at concentrations up to 512 μM (2.6% inhibition at 256 μM and 1.9% inhibition at 512 μM). We also measured the effect of the compound on the voltage-gated cardiac sodium ion channel Nav 1.5. This channel is a key component for the initiation and transmission of the electrical signal throughout the heart. The IC_50_ of NOSO-502 in the patch clamp Nav 1.5 sodium channel assay was higher than 512 μM.

The genotoxic potential of NOSO-502 was investigated using the micronucleus (MN) assay. This test detects both aneugenic (whole chromosome) and clastogenic (chromosome breakage) damage in interphase cells (7). There was no significant increase of micronuclei in cells treated with 512 μM of NOSO-502 *versus* an S9 medium negative control (0.61% cells with micronuclei for NOSO-502 *versus* 0.7% for S9 medium).

NOSO-502 had no cytotoxic effect against mammalian HepG2 (Human hepatocellular carcinoma) cells at concentrations up to 512 μM (0% inhibition from 16 to 256 μM and 4.2% inhibition at 512 μM) and did not show any hemolytic activity at 100 μM. The compound (10 μ M) had no significant activity against any of the 55 cell surface receptors or enzymes tested in a broad-based screen.

### NOSO-502 is resistant to biotransformation by hepatocytes and microsomes

NOSO-502 was resistant to biotransformation when incubated in mouse, rat, dog, monkey, and human liver microsomes and hepatocytes during the *in vitro* study conducted to evaluate metabolic stability. After 45 minutes, 70.5 to 78.6% of NOSO-502 remained after incubation with microsomes of the different species. The half-lives of NOSO-502 in liver microsomes were 116, 129, 101, 147, and 145 min for mouse, rat, dog, monkey, and human, respectively. After 60 minutes, 79.5 to 91.9% of NOSO-502 remained after incubation with hepatocytes of the different species. The half-lives of NOSO-502 in hepatocytes were 192, 194, 483, 698, and 329 min for mouse, rat, dog, monkey, and human microsomes respectively.

### NOSO-502 shows variable stability in plasma of different species

NOSO-502 showed variable stability to biotransformation when incubated in mouse, rat, dog, monkey, and human plasma; 10.1 to 61.2% of NOSO-502 remained after incubation with plasma of the different species over the 120-min test period. The half-lives of NOSO-502 were 54, 36, 158, 96, and 79 min for mouse, rat, dog, monkey, and human plasma, respectively.

### Pharmacokinetics

The pharmacokinetics of NOSO-502 was evaluated in normal female CD-1 mice or normal female Sprague-Dawley rats. NOSO-502 was administered intravenously at 30 mg/kg to mice and 15 mg/kg to rats. The concentration-*versus*-time curves and the results of the pharmacokinetic analysis are summarized in Figure 3. In mice, NOSO-502 displayed moderate clearance (1.13 L/h/kg), a moderate volume of distribution (0.66 L/kg), and a half-life of 25 min. The pharmacokinetics of NOSO-502 in rats showed a longer half-life (90 min) but were consistent with the results in mice, with a plasma clearance of 1.92 L/h/kg and a volume of distribution of 0.94 L/kg. NOSO-502 showed moderate plasma protein binding, with 19.8, 20.5, 17.6, and 18.7% unbound in mouse, rat, dog, and human plasma, respectively.

**Figure 3.**
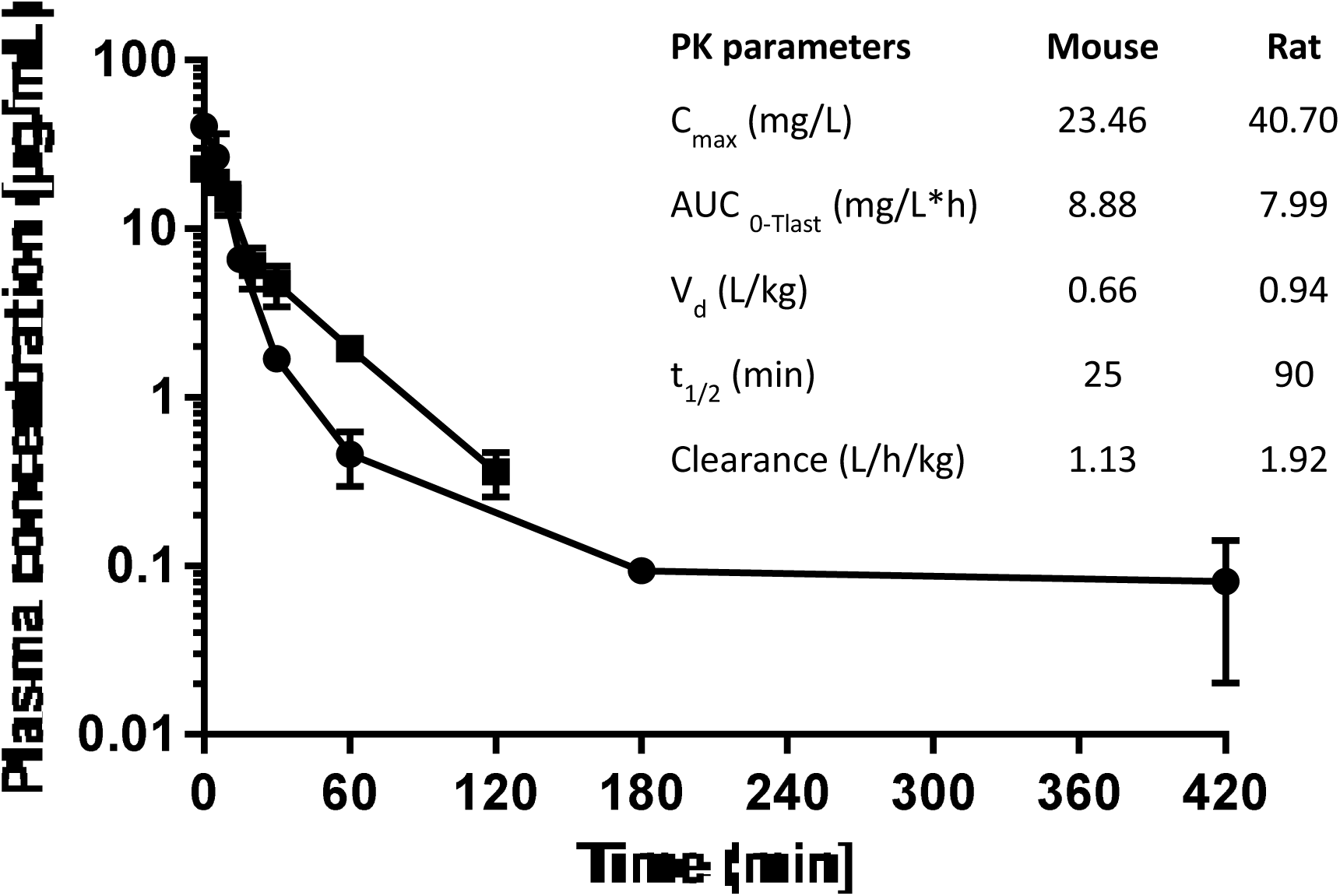
Pharmacokinetic studies with CD-1 mice (closed squares) and SD rats (closed circles) following intravenous dosing at 30 and 15 mg/kg respectively. Each symbol represents the mean and error bars indicate the standard error of the mean.

### NOSO-502 shows efficacy in several murine infection models

The efficacy of NOSO-502 was evaluated in murine infection models to determine whether NOSO-502 has potential as a clinical therapy. *In vivo* efficacy studies were conducted by administering NOSO-502 subcutaneously. The efficacy of NOSO-502 was first assessed in a neutropenic murine sepsis infection model. This model, with *E. coli* EN122 (ESBL, clinical isolate), was established in female NMRI mice. NOSO-502 was administered subcutaneously 1 h post-inoculation at set concentrations of 1.3, 2.5, 5, 10, 20, and 40 mg/kg, whereas colistin was administered by the same route at 5 mg/kg. Five hours post-challenge, blood samples were collected, and the mice euthanized. Blood was serially plated and colonies enumerated to determine the CFU/ml of blood. NOSO-502 was highly effective, achieving an ED_50_ of 3.5 mg/kg and 1-, 2- and 3-log reductions in blood burden at 2.6, 3.8, and 5.9 mg/kg, respectively (Fig. 4).

**Figure 4.**
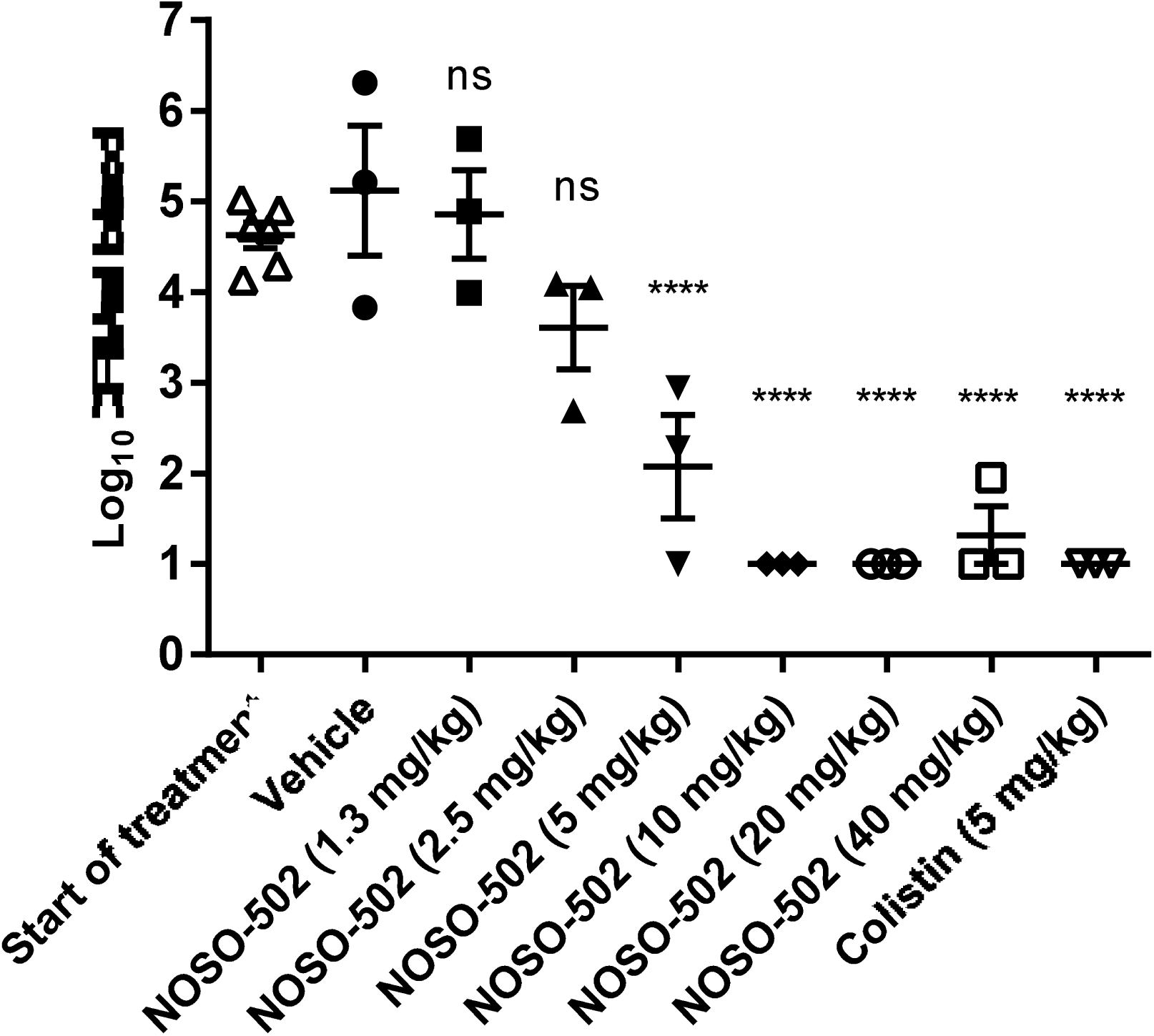
Efficacy of NOSO-502 and colistin in a neutropenic murine sepsis infection model *against E. coli* EN122. Each symbol represents an individual mouse and the horizontal line indicates the mean. Error bars indicate the standard error of the mean. Statistically significant reduction *versus* vehicle control (One-way ANOVA, Dunnett’s comparison): ns, not significant; *, p ≤ 0.05; **, p ≤ 0.01; ***, p ≤ 0.001; ****, p ≤ 0.0001.

**Figure 5.**
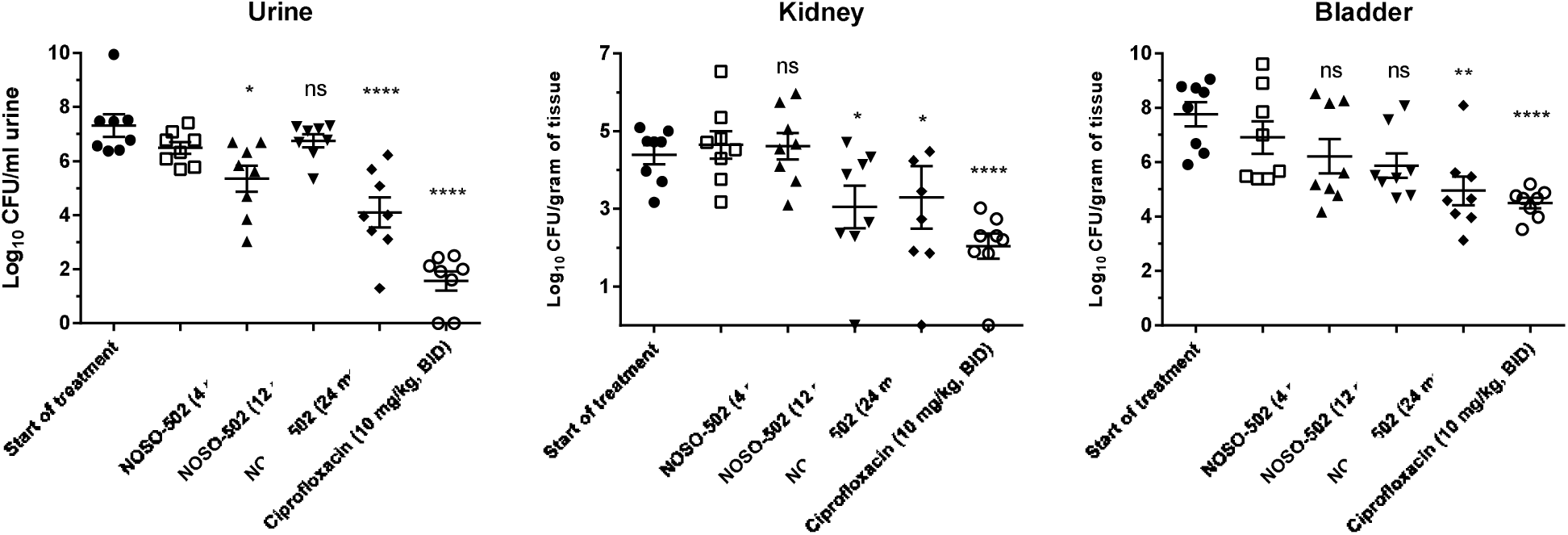
Efficacy of NOSO-502 and ciprofloxacin in a murine UTI model against *E. coli* UTI89. Each symbol represents an individual mouse and the horizontal line indicates the mean. Error bars indicate the standard error of the mean. Statistically significant reduction *versus* vehicle control (Kruskal-Wallis statistical test, multiple comparison): ns, not significant; *, p ≤ 0.05; **, p ≤ 0.01; ***, p ≤ 0.001; ****, p ≤ 0.0001.

A mouse *E. coli* UTI89 upper urinary tract infection model was established in female C3H/HeN mice. Administration of 24 mg/kg NOSO-502 once daily resulted in a statistically significant reduction in urine, bladder, and kidney burdens relative to vehicle control animals. At four days post-infection, NOSO-502 reduced the urine burden by 2.39 log_10_ CFU/ml (P ≤ 0.0001), the bladder burden by 1.96 log_10_ CFU/ml (P = 0.0012), and the kidney burden by 1.36 log_10_ CFU/ml (P = 0.0123) relative to vehicle (**Fig.5**).

A neutropenic mouse *E. coli* ATCC BAA-2469 (NDM-1) intraperitoneal (IP) sepsis infection model was established in male CD-1/ICR mice. Ninety percent of the vehicle-treated mice succumbed to infection prior to the end of the study. All NOSO-502-treated mice (4, 12, and 24 mg/kg) survived up to the end of the study at 24 h (P = 0.0009 relative to vehicle). The vehicle group had a mean and median survival time of 19.8 h and 20.2 h, respectively. One subcutaneous administration of NOSO-502 resulted in statistically significant dose-dependent reductions in blood and IP wash burdens relative to vehicle control animals at all doses. Treatment with 4 mg/kg of NOSO-502 reduced the blood and IP wash burden by 1.48 log_10_ CFU/ml (P = 0.0081) and 0.68 log_10_ CFU/ml (P = 0.0145), respectively. Treatment with 12 mg/kg reduced the blood burden by 2.14 log_10_ CFU/ml (P < 0.0001) and the IP wash burden by 2.07 log_10_ CFU/ml (P ≤ 0.0001) and treatment with 24 mg/kg reduced the blood burden by 2.37 log_10_ CFU/ml (P ≤ 0.0001) and the IP wash burden by 2.74 log_10_ CFU/ml (P ≤ 0.0001) (Fig. 6).

**Figure 6.**
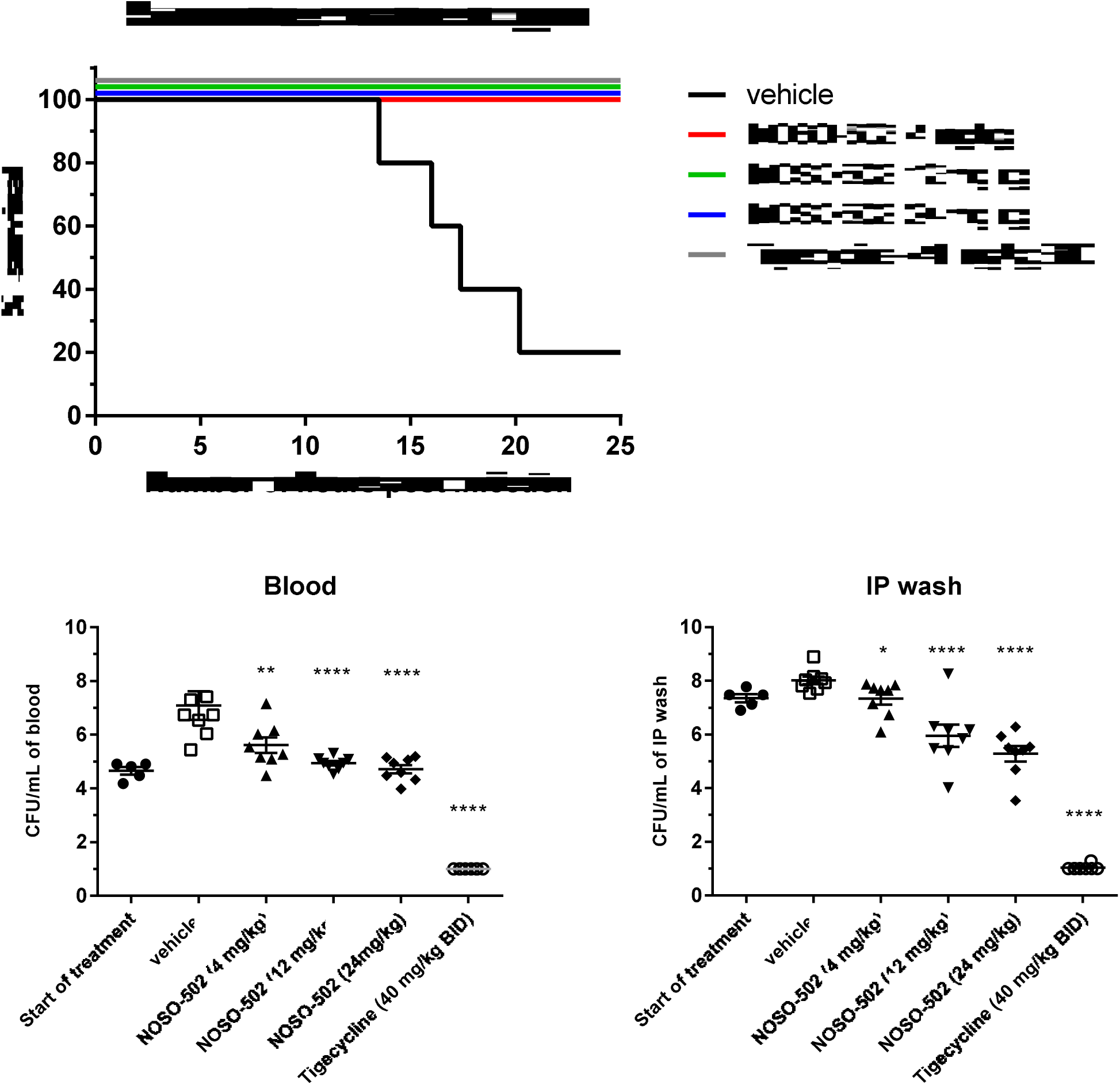
Efficacy of NOSO-502 and tigecycline in a survival neutropenic sepsis infection model against *E. coli* ATCC BAA-2469 (NDM-1). Each symbol represents an individual mouse and the horizontal line indicates the mean. Error bars indicate the standard error of the mean. Statistically significant reduction *versus* vehicle control (Kruskal-Wallis statistical test, multiple comparison): ns, not significant; *, p ≤ 0.05; **, p ≤ 0.01; ***, p ≤ 0.001; ****, p ≤ 0.0001.

A neutropenic mouse *K. pneumoniae* NCTC 13442 (OXA-48) lung infection model was established in male CD-1/ICR mice. NOSO-502 was administered subcutaneously 2 h, 8 h, 14 h and 20 h post-inoculation at set concentrations of 2, 6, and 20 mg/kg (equivalent to 8, 24 and 80 mg/kg/day), whereas tigecycline was administered by the same route at 40 mg/kg (equivalent to 160 mg/kg/day). NOSO-502 was also administered once 2 h post-inoculation at 80 mg/kg. Twenty-six hours post-challenge, mice were euthanized, and the lungs collected. Administration of NOSO-502 resulted in statistically significant reductions in lung burdens relative to vehicle control animals at all doses. Treatment with 8, 24 and 80 mg/kg/day of NOSO-502 reduced the lung burden by 2.69, 3.99 and 4.07 log_10_ CFU/gram of lung tissue respectively (P ≤ 0.0001). Treatment with 80 mg/kg once reduced the lung burden by 3.98 log_10_ CFU/ gram of lung tissue (P ≤ 0.0001) and treatment with 160 mg/kg/day of tigecycline reduced the lung burden by 3.14 log_10_ CFU/ gram of lung tissue (P ≤ 0.0001) (Fig. 7).

**Figure 7.**
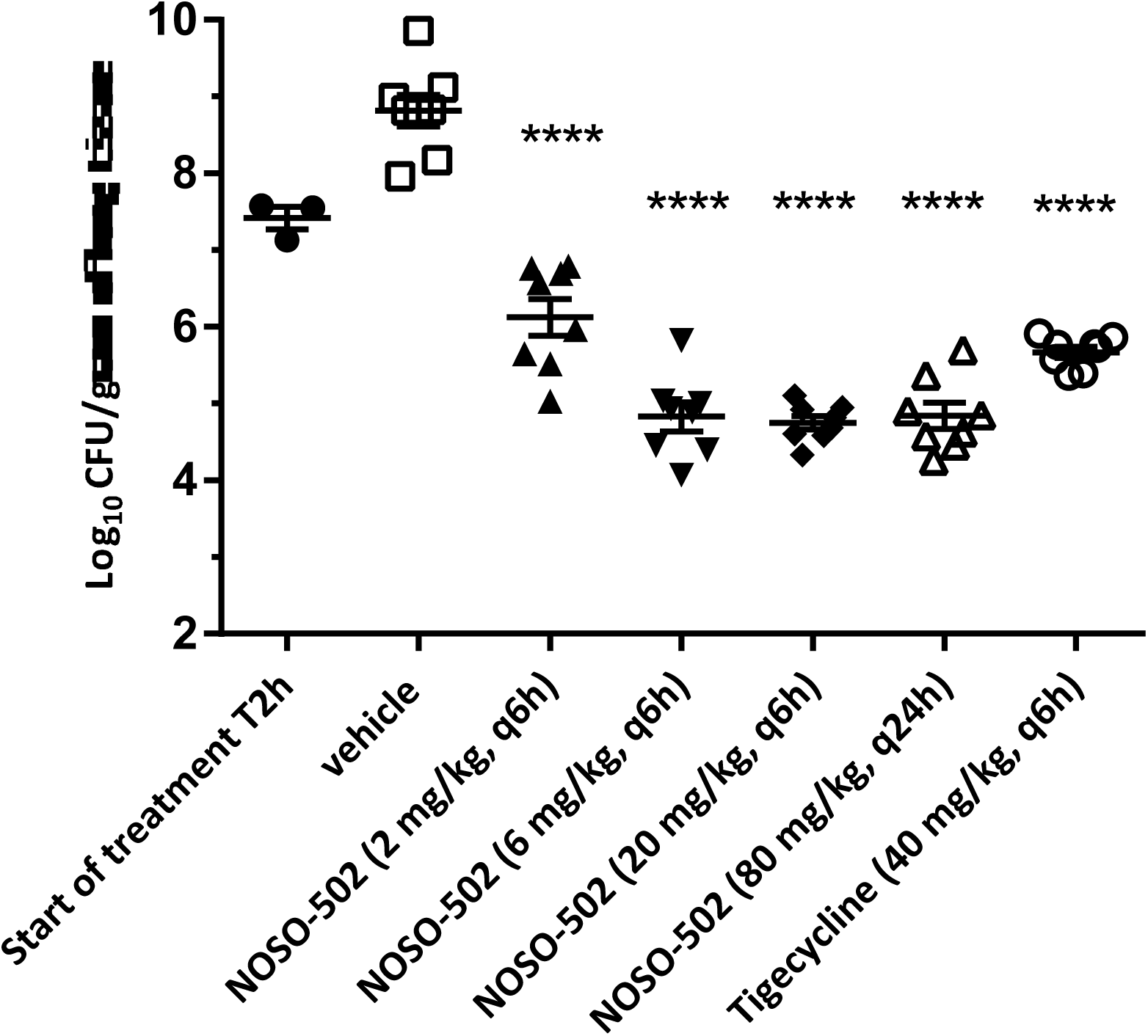
Efficacy of NOSO-502 and tigecycline in a murine lung infection model against *K. pneumoniae* NCTC 13442 (OXA-48). Each symbol represents an individual mouse and the horizontal line indicates the mean. Error bars indicate the standard error of the mean. Statistically significant reduction *versus* vehicle control (One-way ANOVA, Dunnett’s comparison): ns, not significant; *, p ≤ 0.05; **, p ≤ 0.01; ***, p ≤ 0.001; ****, p ≤ 0.0001.

## DISCUSSION

The urgent need to discover new antibiotics active against Gram-negative bacteria with a novel mechanism of action to counter the threat of drug-resistant infection is widely recognized. NOSO-502 is the first preclinical candidate of a novel antibiotic class, the Odilorhabdins (ODLs). ODLs are cationic peptides that inhibit bacterial translation by a novel mechanism of action. ODLs bind to the small subunit of bacterial ribosomes at a site not exploited by any known ribosome-targeting antibiotic. When bound to the ribosome, ODLs make contacts with both the rRNA and tRNA and kill bacteria by interfering with the decoding of genetic information and inhibiting ribosome progression along the mRNA in a context-specific manner (3).

NOSO-502 is active against *Enterobacteriaceae*, including CRE belonging to all classes of the Ambler classification and resistant to gentamicin, polymyxin B, or tigecycline. This is crucial, because these antibiotics, classically used for the treatment of such infections, are associated with high levels of resistance ranging from 9.7 to 51.3% (mean 22.6%) for colistin, 5.6 to 85.4% (mean 43.5%) for gentamicin, and 0 to 33% (mean 15.2%) for tigecycline (8, 9, 10, 11, 12, 13, 14, 15, 16, 17). Current options to address these resistance issues are not entirely satisfactory, because none of the recently approved antibiotics or those under development are effective against all CRE. The combination Ceftazidime-avibactam displays *in vitro* activity against CRE isolates that produce KPC, AmpC and OXA enzymes. However, this drug is not active against metallo-β-lactamases, such as NDM, IMP, or VIM (18). This combination was approved by the US Food and Drug Administration in 2015 and by the European Medicines Agency in 2016 for treating complicated urinary tract and intra-abdominal infections. None of the novel antimicrobials (plazomicin, a new aminoglycoside or eravacycline, a new tetracycline) or novel combinations as aztreonam and avibactam, meropenem and vaborbactam, imipenem and relebactam/cilastatin, or ceftaroline fosamil and avibactam are effective against all classes of carbapenemases like NOSO-502 (19). Recently, CRE have caused numerous outbreaks of severe nosocomial infections and have become endemic in several countries (20, 21, 22, 23, 24). These infections have been associated with mortality rates exceeding 50% in some reports (25, 26, 27, 28). NOSO-502 can overcome multiple mechanisms of colistin-resistance strains. Furthermore, the compound demonstrated rapid bactericidal activity and a low potential for the development of resistance.

NOSO-502 is effective in mouse models of serious hospital-acquired infections. It provided significant protection against the Gram-negative pathogens *E. coli* and *K. pneumoniae*, the highest-incidence pathogens in complicated intra-abdominal and urinary tract infections, in septicemia following peritoneal challenge, and in acute pyelonephritis. NOSO-502 was active in mouse infection models against *E. coli* strains expressing the metallo-β-lactamase NDM-1 and resistant to other major antibiotic classes, including fluoroquinolones, macrolides, aminoglycosides, β-lactams, cephalosporins, and carbapenems. These results are encouraging and show the strong potential for *in vivo* efficacy of NOSO-502. Effective doses will be optimized after the best dosing schedule is defined during a PKPD study.

NOSO-502 showed a good safety profile, with no *in vitro* nephrotoxicity, cardiotoxicity, genotoxicity, or cytotoxicity at concentrations up to 512 μM. Nephrotoxicity is a serious side effect of many drugs, including cationic antibiotics aminoglycosides and polymyxins (29, 30, 31). Polymyxins accumulate extensively within proximal tubular cells (PTCs) of the kidneys, where they induce damage, which may lead to acute kidney injury (AKI) in patients (32). AKI is the major dose-limiting adverse effect of this class of antibiotics and affects 50 to 60% of patients receiving them (31, 33). Aminoglycosides are filtered across the glomerulus and then excreted, with 5 to 10% of a parenteral dose being taken up and sequestered by the PTCs, in which the aminoglycoside can achieve high concentrations (34). AKI due to acute tubular necrosis is a relatively common complication of aminoglycoside therapy and affects 10 to 20% of patients (29, 30). The results of NOSO-502 on HRPTEpiC and HK-2 cells are promising, but must be confirmed by histopathological examination of kidney cells following *in vivo* administration to animals, the standard assay for studying nephrotoxicity effects.

Cardiotoxicity issues are associated with many antibiotics, including macrolides, ketolides, and fluoroquinolones. These classes have been associated with prolongation of cardiac repolarization. All these agents produce a blockage of the hERG channel-dependent potassium current in myocyte membranes, resulting in a prolonged QTc interval which may give rise to ventricular fibrillation or tachycardia (35). Nav 1.5 is another channel involved in cardiotoxicity issues. Its activation induces depolarization of the cell membrane. Failure of the Nav 1.5 sodium channel to adequately conduct the electrical current across the cell membrane can result in a potentially fatal disorder. NOSO-502 did not show any effects on hERG or Nav 1.5 channels at high concentrations.

Here, we confirmed that NOSO-502, like many other therapeutic peptides, is safe and highly selective. NOSO-502 interacts strongly with a specific site on the 30S subunit of bacterial ribosomes but has no significant activity against any of the 55 cell surface receptors, transporters, or ion channels tested. There is increasing interest in peptides in pharmaceutical research and development (R&D) and approximately 140 are currently being evaluated in clinical trials and more than 500 are in preclinical development (36, 37). The main limitation of peptides is their predisposition to enzymatic degradation. Thus, most do not circulate in blood for more than a few minutes, preventing their usefulness as therapeutic agents. On the contrary, NOSO-502 showed good stability in plasma, microsomes, and hepatocytes, probably due to the presence in its structure of three non-standard amino-acid residues: α, γ-diamino-β-hydroxy butyric acid (Dab(β OH)) at position 2 (*N*-terminus), α, β –dehydro arginine (Dha) at position9 (C-terminus), and D-ornithine at position 5. This translates into relatively long half-lives in mice and rats.

NOSO-502 represents a new class of very promising bacterial ribosomal inhibitors to combat bacterial multidrug resistance.

## MATERIALS AND METHODS

### Bacterial strains and antimicrobial agents

Reference strains are from the German collection of microorganism and cell cultures (DSM), the American Type Culture Collection (ATCC), the National Collection of Type Cultures (NCTC), and the Medical and Molecular Microbiology, Faculty of Science and Medicine, University of Fribourg, Switzerland. Clinical strains used to determine the MIC_90_ of NOSO-502 come from Warsaw, Copenhagen, Cardiff, and Madrid hospitals. NOSO-502 was synthesized at Nosopharm, Nîmes, France. Ciprofloxacin (Sigma-Aldrich, ref: 1134335), gentamicin (Sigma-Aldrich, ref: G1397), imipenem (Sigma-Aldrich, ref IO160), polymyxin B (Sigma-Aldrich, ref: 92283), and tigecycline (Sigma-Aldrich, ref: PZ0021) were provided by the manufacturers as standard powders except for gentamicin and polymyxin B, in solution at 50 and 20 mg/ml respectively.

### Minimum inhibitory concentration (MIC)

MIC values were determined using Clinical and Laboratory Standards Institute (CLSI) broth microdilution methodology, colony direct suspension, as described in CLSI document M07-A10 (38).

### Time-dependent killing

Time-kill assays were performed by the broth macrodilution method, according to the CLSI guidelines M26-A (39).

For preparing the inoculum, between 5 and 30 colonies of a single morphological type from a 16- to 24-h Mueller–Hinton agar plate (MHA) were picked and used to inoculate a tube containing 5 ml prewarmed cation-adjusted Mueller–Hinton broth (CA-MHB). The bacterial suspension was incubated at 35°C until it was visibly turbid. The turbidity of the actively growing broth culture was adjusted with CA-MHB to obtain a calculated OD_600_ between 0.11 and 0.15. Shaken flasks (250 ml) containing 50 ml CA-MHB, with the appropriate NOSO-502 concentrations, were inoculated with 0.5 ml exponentially grown bacteria suspension (5 × 10^7^ cells/ml) to yield a final concentration of approximately 5 × 10^5^ cells/ml. Two multiples of the MICs (four and eight) were used to detect differences in killing. Flasks were incubated at 35°C, with shaking at 150 rpm, and aliquots removed at 0, 1, 2, 3, 4, 6, and 24 h for the determination of viable counts. Serial dilutions were prepared in a sterile 0.9% sodium chloride solution and plated on MHA plates. The plates were incubated at 35°C for 24 h, and the number of colonies determined. The detection level by this plating method was 50 CFU/ml. Killing curves were constructed by plotting the log_10_ CFU/ml *versus* time over 24 h and the change in bacterial concentration determined.

### Determination of mutation frequency

Bacterial strains were grown in antibiotic-free Luria Bertani broth at 35°C for 18 h. Approximately 10^9^ CFU of each strain were plated in duplicate onto MHA plates containing NOSO-502 concentrations at 4× and 8× the MIC values. The plates were read after 24 and 48 h of incubation at 35°C. The frequency of selected resistant mutants was calculated as the ratio of the number of bacteria growing divided by the number of bacteria in the original inoculum, which was calculated by plating several dilutions of the original inoculum.

### Multiplexed HRPTEpiC cytotoxicity assay

The multiplexed cytotoxicity assay on human renal proximal tubule epithelial cells (HRPTEpiC) was conducted by Eurofins Panlabs (Eurofins Panlabs, Inc. St Charles, MO, USA) by using an image-based High Content Analysis (HCA) technique where cells were fixed and stained with nuclear dye to visualize nuclei and fluorescently labeled antibodies to detect drug induced cellular injury and cellular stress arising from oxidative and chemical stress. Cells were seeded into 384-well plates and grown in RPMI1640, 10% FBS, 2 mM L-alanyl-L-Glutamine, 1 mM Sodium Pyruvate in a humidified atmosphere of 5% CO_2_ at 37°C. NOSO-502, gentamicin and polymyxin B were added 24 h post cell seeding. Compounds were serially diluted 3.16-fold and assayed over 10 concentrations in a final assay concentration of 0.5% DMSO from 100 μM to 3.7 nM. At the same time, a time zero untreated cell plate was generated. After a 48-h incubation period, cells were fixed and stained with fluorescently labeled antibodies and nuclear dyed to allow visualization of nuclei, injured cells and stressed cells. Injured cells were detected using a KIM-1 (Kidney Injury Molecule-1) antibody. Stressed cells were detected using an anti-HSP27 (Heat Shock Protein 27) antibody. Cell proliferation was measured by the signal intensity of the incorporated nuclear dye. The cell proliferation assay output was referred to as the relative cell count. To determine the cell proliferation end point, the cell proliferation data output was transformed to percent of control (POC) using the following formula: POC = relative cell count (compound wells) / relative cell count (vehicle wells) × 100. The signal intensity of the incorporated cellular stress and injury measurements were normalized with the relative cell count from each well. Automated fluorescence microscopy was carried out using a Molecular Devices ImageXpress Micro imager, and images were collected with a 4× objective.

### Cytotoxicity testing

HK-2 and HepG2 cytotoxicity assays were run by Eurofins-Cerep (Cerep Cytotoxicity Profile, Eurofins-Cerep SA, Poitiers, France) as described in reference (40). Cell viability was measured using a luciferase-coupled ATP-quantitation assay (CellTiter-Glo; Promega, Madison, WI, USA). In this assay, luminescent signal is proportional to the amount of ATP and thus to the number of metabolically competent cells; cell injury and death result in a marked decrease in intracellular ATP levels. HK-2 and HepG2 cells were dispensed at 6,000– 3,000 cells/5 μl/well in white tissue-culture treated 96-well solid-bottom assay plates and incubated at 37°C for 16 h, to allow cell attachment, followed by the addition of NOSO-502 at 16, 32, 64, 128, 256, and 512 μM. After compound addition, plates were incubated for 48 h at 37°C. At the end of the incubation period, 5 μl CellTiter-Glo reagent was added, the plates were incubated at room temperature for 30 min, and the luminescence intensity of each well was determined. Each experiment was carried out in duplicate and the results are reported as the average percent of cytotoxicity for each test concentration and as IC_50_ value (concentration producing a half-maximal inhibition of control response = half maximal cytotoxicity) determined by non-linear regression analysis of the concentration-response curve generated with mean replicate values using Hill equation curve fitting.

### hERG tail current inhibition

Inhibition of the human ether-a-go-go-related gene (hERG) cardiac potassium ion channel was determined by Eurofins Panlabs (, St Charles, MO, USA) in CHO-K1 (Chinese Hamster Ovary) cells stably transfected with human hERG cDNA using QPatch Automated whole-cell patch clamp electrophysiology as described in reference (41). NOSO-502 was tested at 64, 256 and 512 μM, the extracellular solution (control) is applied first and the cell is stabilized in the solution for 5 min. Then the test compound is applied from low to high concentrations sequentially on the same cell with 5min each test concentration at room temperature.

### Nav 1.5 peak current inhibition

Inhibition of the Nav 1.5 human sodium ion channel was determined by Eurofins Panlabs (, St Charles, MO, USA) in HEK-293 cells stably transfected with human Nav1.5 cDNA (type V voltage-gated sodium channel alpha subunit, accession #NM_000335) using IonWorks Quattro Automated whole-cell patch clamp electrophysiology. NOSO-502 was tested at 4, 8, 16, 32, 64, 128, 256 and 512 μM, the voltage protocol is applied prior to compound addition (Pre), the compounds are added and incubated for 600 seconds at room temprature, and then the voltage protocol is applied a final time (Post) on the IonWorks Quattro.

### *In vitro* Micronucleus assay

The test was conducted by Eurofins Panlabs (, St Charles, MO, USA).

CHO-K1 cells were pre-loaded with a cell dye that stains the cytoplasm, after which the cells were treated with NOSO-502 at 32, 64, 128, 256 and 512 μM for 24 h. At the end of the incubation period the cells were fixed, and their DNA was stained with Hoechst. The visualization and scoring of the cells was done using an automated fluorescent microscope coupled with proprietary automated image analysis software (42). The percent of micronucleated cells is calculated. A marginally-positive result (“-/+”) is defined as a value significantly higher than controls (t-test, p < 0.05), and at least 2-fold higher than controls. A positive result (“+”) is defined as a value significantly higher than controls (t-test, p < 0.05) and at least 3-fold higher than controls.

### Hemolytic activity

Mouse red blood cells were washed with Phosphate Buffered Saline (PBS) until the supernatant was clear after centrifugation and resuspended in PBS to 10% (v/v). Three hundred microliters of the suspension were added to a microtube containing an equal volume of NOSO-502 to give a final concentration of 100 μM. PBS and deionized water were used as 0 and 100% hemolytic controls, respectively. Plates were incubated at 35°C for 45 min. Subsequently, the microtube was centrifuged and the supernatant transferred to a new microtube. The release of hemoglobin in the supernatant was monitored by absorbance at 540 nm. Experiments were performed in triplicates.

### Hepatocyte stability

The hepatocyte metabolic stability assays were performed by Cyprotex discovery Ltd. (Macclesfield, UK). This assay utilizes cryopreserved pooled hepatocytes from different species (mouse, rat, dog, monkey and human), stored in liquid nitrogen prior to use. Williams E media supplemented with 2 mM L-glutamine, 25 mM HEPES and NOSO-502 (NOSO-502 final substrate concentration 1 μM, test compound prepared in water; control compound final substrate concentration 3 μM, final DMSO concentration 0.25%) were pre-incubated at 37°C prior to the addition of a suspension of cryopreserved hepatocytes (final cell density 0.5 ×10^6^ viable cells/ml in Williams E media supplemented with 2 mM L-glutamine and 25 mM HEPES) to initiate the reaction. The final incubation volume was 500 μl. Two control compounds were included with each species alongside an appropriate vehicle control. The reactions were stopped by transferring an aliquot of the mixture to 40% trichloroacetic acid (TCA) in water, containing internal standard for the test compounds (NOSO-95216, 1 μM final concentration), or methanol for the control compounds, at various time points (0, 5, 10, 20, 40 and 60 min). The termination plates were centrifuged at 2,500 rpm at 4°C for 30 min to precipitate the protein. Following protein precipitation, the test compound sample supernatants were diluted with analytical grade water, whereas the control compounds were diluted with internal standard (metoprolol) in water. The test compound samples were analyzed by LC-MS/MS. The gradient of the line was determined from a plot of ln peak area ratios (compound peak area/internal standard peak area) against time. Subsequently, half-life and intrinsic clearance were calculated using the following equations: elimination rate constant (k) = (- gradient); half-life (t½) (min) = 0.693/k; intrinsic clearance (CLint) (μl/min/million cells) = (V x 0.693)/t½ where V = incubation volume (μl)/ Number of cells. Relevant control compounds were assessed, ensuring that intrinsic clearance values fell within the specified limits (if available).

### Microsome stability

The microsome metabolic stability assays were performed by Cyprotex discovery Ltd. (Macclesfield, UK). Pooled microsomes from different species (mouse, rat, dog, monkey, and human) were stored at -80°C prior to use. Microsomes (final protein concentration mg/ml), 0.1 M phosphate buffer pH 7.4, and NOSO-502 (test compound final substrate concentration 1 μM, test compound prepared in water; control compound final substrate concentration 3 μM, final DMSO concentration 0.25%) were pre-incubated at 37°C prior to the addition of NADPH (final concentration 1 mM) to initiate the reaction. The final incubation volume was 500 μl. A minus cofactor control incubation was included for each compound tested, in which 0.1 M phosphate buffer pH 7.4 was added instead of NADPH (minus NADPH). Two control compounds were included with each species. Each compound was incubated for 0, 5, 15, 30 and 45 min. The control (minus NADPH) was incubated for 45 min only. The reactions were stopped by transferring an aliquot of the mixture to 40% TCA in water containing internal standard (NOSO-95216, 1 μM final concentration) for the test compounds, or methanol for the control compounds, at the indicated time points. The termination plates were centrifuged at 2,500 rpm for 20 min at 4°C to precipitate the protein. Following protein precipitation, the test compound sample supernatants were diluted with analytical grade water, whereas the control compounds were diluted with internal standard (metoprolol) in water. The test compound samples were analyzed by LC-MS/MS. The gradient of the line was determined from a plot of ln peak area ratios (compound peak area/internal standard peak area) against time. Subsequently, the half-life and intrinsic clearance were calculated using the following equations: elimination rate constant (k) = (- gradient); half-life (t^1/2^) (min) =0.693/k; intrinsic clearance (CLint) (μl/min/mg protein) = (V × 0.693)/t^1/2^ where V = incubation volume (μl)/microsomal protein (mg). Relevant control compounds were assessed, ensuring that intrinsic clearance values fell within the specified limits (if available).

### Plasma stability

The plasma stability assays were performed by Cyprotex Discovery Ltd. (Macclesfield, UK). Species-specific plasma (heparin anti-coagulant) was adjusted to pH 7.4 at 37°C and NOSO-502 or control compound (test compound final substrate concentration 10 μM, test compound prepared in water; control compound final substrate concentration 1 μM, final DMSO concentration 2.5%) was added to initiate the reaction. The final incubation volume was 200 μl. All incubations were performed singularly for each compound at each time point. A vehicle control incubation was included using either water or DMSO, along with a control compound known to be metabolized specifically by each species. Each compound was incubated for 0, 15, 30, 60, and 120 min at 37°C. The reactions were stopped by the addition of 40% TCA in water containing internal standard (NOSO-95216, 1 μM final concentration) for the test compounds, or methanol for the control compounds, at the appropriate time points. The vehicle control incubation was incubated for 120 min only. The termination plates were centrifuged at 3,000 rpm for 45 min at 4°C to precipitate the protein. Following protein precipitation, the test compound sample supernatants were diluted with analytical grade water, whereas the control compounds were diluted with internal standard (metoprolol) in water. The test compound samples were analyzed by LC-MS/MS. The percentage of parent compound remaining at each time point relative to the 0 min sample was calculated from peak area ratios (compound peak area/internal standard peak area).

### Selectivity profile

The affinity of NOSO-502 tested at 10 μM was assessed using radioligand binding assays for 55 cell surface receptors, transporters, and ion channels were tested by Eurofins-Cerep (Cerep Diversity Profile, Eurofins-Cerep SA, Poitiers, France). Receptors tested included those to adenosine (A_1_, A_2A_, A_3_), adrenergic (alpha^1^, alpha_2_, beta_1_, beta_2_), angiotensin-II (AT_1_), bradykinin (B_2_), cannabinoid (CB_1_), chemokines (CCCR_1_, CXCR_2_) cholecystokinin (CCK_1_), dopamine (D_1_, D_2S_), endothelin (ET_a_), GABA non-selective, galanine, (GAL_2_), histamine (H_1_, H_2_), melanocortin (MC_4_), muscarinic (M_1_, M_2_, M_3_), neurokinin (NK_2_, NK_3_), neuropeptide Y (Y_1_, Y_2_), neurotensin (NTS_1_), opioid and opioid-like (delta_2_, kappa, mu, NOP), prostanoid (EP_4_), serotonin (5-HT_1A_, 5-HT_1B_, 5-HT_2A_, 5-HT_2B_, 5-HT_5A_, 5-HT_6_, 5-HT_7_), somatostatin (sst), vasoactive intestinal peptide (VPAC_1_), and vasopressin (V_1a_). Transporters tested included the dopamine transporter (DAT), norepinephrine transporter (NET), and serotonin transporter (5-HT). ion channels tested included those for potassium (K_v_ and SK_ca_ channels) calcium (Ca^2^+ channel, L-type, verapamil site), sodium (Na^+^ channel site 2), GABA (BZD and Cl^-^ channel GABA gated), and serotonin (5- HT_3_). Receptor, transporter, or ion channel binding by a specific ligand was defined as the difference between total and nonspecific binding, determined in the presence of an excess of unlabelled ligand. Results are expressed as the percent inhibition of control-specific binding or the percent variation of control values obtained in the presence of NOSO-502.

### Pharmacokinetic analysis

Pharmacokinetics were performed by Pharmacelsus (Saarbrücken, Germany). CD-1 female mice and female Sprague Dawley rats were injected intravenously with 30 and 15 mg/kg of NOSO-502, respectively, prepared in saline (5 ml/kg). Blood (100 μl) was collected from three animals per time point (5, 10, 20, 30, 60 and 120 min post-dose for mice and 5, 15, 30, 60, 180 and 420 min post-dose for rats) in tubes containing K3-EDTA as anticoagulant. Samples were stored on ice and centrifuged at 6,000 rpm for 10 min at 4°C. A sample volume of 50 μl was mixed with 5 μl solvent (acetonitrile/H_2_O/DMSO, (5/4/1, v/v/v) + 1% formic acid). A volume of 10 μl solvent containing the internal standard and 50 μl precipitant (10% TCA) were added to 55 μl of sample. The mixture was vortexed and centrifuged at 6,000 x g (room temperature) for 10 min. The protein-free supernatant was analysed by LC-MS using an Ultimate 3000RS U-HPLC coupled with an Orbitrap Q Exactive mass spectrometer.

Analytes were separated on a Accurore phenyl-hexyl analytical column (2.1 × 50 mm, 2.6 μM, Thermo, Germany) using a linear gradient of mobile phase A (acetonitrile/0.2% heptafluorobutyric acid)-mobile phase B (water/0.2% heptafluorobutyric acid), starting from 5% of mobile phase A to 97% in 2.2 min, and a flow rate of 0.6 μl/min. Peaks were analysed by mass spectrometry (ESI ionization in MRM mode) using Xcalibur 4.0 software. The products [M+2H]^2+^ and [M+3H]^3+^ analysed were 539.8 and 360.2 Da, respectively. PK parameters were calculated using a non-compartmental analysis model and Kinetica 5.0 software (Thermo Scientific, Waltham, USA). The mean plasma concentrations from all three mice at each time point were used in the calculation.

### Plasma protein binding

The plasma stability assays were performed by Cyprotex Discovery Ltd. (Macclesfield, UK). This study was conducted to determine the extent of binding of NOSO-502 to the proteins in human, monkey, dog, rat and mouse plasma. Solutions of NOSO-502 or control compound (NOSO-502 final substrate concentration 2 μM in water; control compound final substrate concentration 2 μM, final DMSO concentration 0.5 %) were prepared in 100% species-specific plasma (collected using EDTA as the anti-coagulant). The experiment was performed using equilibrium dialysis (RED device) with the two compartments separated by a semi-permeable membrane. Buffer (pH 7.4) was added to one side of the membrane and the plasma solution to the other. After equilibration (4 h), samples were taken from both sides of the membrane. Calibration standards were prepared in plasma and buffer. All incubations were performed in triplicate. A control compound was included in each experiment. Incubation of the control compound samples was terminated with acetonitrile containing internal standard (metoprolol). Incubation of the test compound samples was terminated with 40% TCA in water containing internal standard (NOSO-95216, 1 μM final concentration). All samples were centrifuged and further diluted with water prior to analysis. The solutions for each batch of control compounds were combined into two groups (protein-free and protein-containing) and cassette analysed by LC-MS/MS using two sets of calibration standards for protein-free (seven points) and protein-containing solutions (seven points).

### **Mouse neutropenic peritonitis/sepsis model**. NOSO-502 was tested against *E. coli EN122* (MIC

= 4 μg/ml, ESBL, clinical isolate 106-EC-09, Denmark) in a murine neutropenic peritonitis/sepsis model. Female NMRI mice (Taconic Biosciences A/S, Lille Skensved, Denmark) were used. Mice had *ad libitum* access to domestic quality drinking water and food (2016 16% Protein Rodents Diet, Harlan, USA) and were exposed to a 12-h light/dark cycle. All animal experiments were approved by the National Committee of Animal Ethics, Denmark, and adhered to the standards of EU Directive 2010/63/EU. Mice were allowed to acclimatize for four days and thereafter neutropenia was induced by intraperitoneal injections of cyclophosphamide (Baxter A/S Søborg Denmark) four days (200 mg/kg) and one day (100 mg/kg) prior to inoculation. Overnight *E. coli* colonies were suspended in saline to 10^7^ CFU/ml and mice were inoculated intraperitoneally with 0.5 ml of the suspension. At 1 h post-inoculation, mice were treated with 1.3, 2.5, 4, 10, 20, or 40 mg/kg NOSO-502, vehicle (PBS pH 7.4) or 5 mg/kg colistin (Sigma-Aldrich, ref: 4461), subcutaneously as a single dose in 0.2 ml. Four hours after treatment, mice were anesthetized, and blood was collected by axillary cutdown. Blood samples were serially diluted and plated on blood agar plates (SSI Diagnostica, Hillerød, Denmark) with subsequent counting of colonies after incubation overnight at 35° C in ambient air. The mice were observed during the study for clinical signs of infection, such as lack of curiosity, social withdrawal, changes in body position and patterns of movement, distress, or pain.

### **Mouse UTI model**. *Ethics statement*

All animal studies were performed under UK Home Office License P2BC7D240 with local ethical committee clearance. All studies were performed by technicians who completed parts A, B, and C of the UK Home Office Personal License course and hold current personal licenses. All experiments were performed in dedicated Biohazard 2 facilities (this site holds a Certificate of Designation).

NOSO-502 was tested against *E. coli* UTI89 (MIC= 4 μg/ml) in a mouse urinary tract infection (UTI) model by Evotec (Manchester, UK). Female C3H/HeN mice 18-22g (Janvier laboratories, UK) were allowed to acclimatize for seven days. Following acclimatization, drinking water was replaced by water containing 5% glucose from five days pre-infection. Previously prepared frozen stocks of *E. coli* UTI89 were diluted to 1 × 10^10^ CFU/ml immediately prior to infection. Mice were infected by directly administering 0.05 ml inoculum (5 × 10^8^ CFU/mouse) *via* the urethra into the bladder under parenteral anesthesia (90 mg/kg ketamine and 9 mg/kg xylazine).

Bladders were emptied prior to infection, and once infected, infection catheters were left in the urinary tract for 10 min to reduce the risk of the organism flowing back out. Following catheter removal mice were allowed to fully recover in warmed humidified cages. Dose formulations of NOSO-502 were prepared in 25 mM PBS. Treatment with 4, 12 and 24 mg/kg NOSO-502 was initiated 24 h post-infection and was administered once daily (q24h) by subcutaneous injection or intravenously (ciprofloxacin) for three days. Mice were euthanized 96 h post-infection (three doses administered). Ciprofloxacin (Bayer, Lot BXHEFTI), administered at 10 mg/kg/dose IV BID, was included as a comparator (six doses administered) and 25 mM PBS was used as vehicle. Urine was collected 24 h post-infection from all animals and used to assess the infection level of each mouse prior to initiation of treatment; all mice were successfully infected. In addition, five mice were euthanized by pentobarbitone overdose to provide a 24 h pretreatment control group. The clinical condition and body weight of all remaining animals were assessed and urine samples collected 96 h post-infection. Animals were then euthanized by pentobarbitone overdose and the kidneys and bladders removed and weighed. Tissue samples were homogenized using a Precellys 24 dual-bead beater in 2 ml ice cold sterile PBS. Homogenates and urine samples were quantitatively cultured onto MacConkey’s agar plates and incubated at 37°C for 24 h before colonies were counted. The data from the culture burdens were analyzed using appropriate non-parametric statistical models (Kruskal-Wallis using Conover-Inman to make all pairwise comparisons between groups) with StatsDirect software v. 2.7.8. and compared to pretreatment and vehicle controls.

### Mouse neutropenic IP sepsis model. *Ethics statement*

All animal studies were performed under UK Home Office License P2BC7D240 with local ethical committee clearance. All studies were performed by technicians who completed parts A, B, and C of the UK Home Office Personal License course and hold current personal licenses. All experiments were performed in dedicated Biohazard 2 facilities (this site holds a Certificate of Designation).

NOSO-502 was tested against *E. coli* ATCC BAA-2469 (MIC= 2 μg/ml) in a IP sepsis model by Evotec (Manchester, UK). Male CD1/ICR mice 25-30g (Charles River, UK) were allowed to acclimatize for 11 days. Mice were rendered neutropenic with two intraperitoneal injections of 150 mg/kg cyclophosphamide four days before infection and 100 mg/kg one day before infection. Previously prepared frozen stocks of *E. coli* ATCC BAA-2469 were diluted immediately prior to infection to 6.8 × 10^7^ CFU/ml. Mice were infected by directly administering 0.5 ml inoculum (3.4 × 10^7^ CFU/mouse) *via* intraperitoneal injection. Dose formulations of NOSO-502 were prepared in 25 mM PBS. Treatment was initiated 1 h post-infection and NOSO-502 doses (4, 12, and 24 mg/kg) were administered once by subcutaneous injection. Tigecycline (MIC = 0.5 μg/mL), administered at 40 mg/kg/dose SC BID, was included as a comparator and two doses were administered. Animals from the pretreatment groups were euthanized 1 h post-infection and all remaining mice were euthanized 25 h post-infection. The clinical condition and body weight of all remaining animals were assessed 25 h post-infection or when animals reached the ethical severity endpoint (whichever came first). Mice were anaesthetized using 2.5% isofluorane/97.5% oxygen followed by a pentobarbitone overdose. When mice were deeply unconscious, blood was collected from all animals under terminal cardiac puncture into EDTA blood tubes. In addition, an intraperitoneal wash with sterile PBS (2 ml IP injected, 1 ml removed for culture) was collected. Five mice were also euthanized by pentobarbitone overdose to provide a 1-h pretreatment control group. Blood and IP wash samples were quantitatively cultured onto CLED agar plates and incubated at 37° C for 24 h before colonies were counted. The data from the culture burdens were analyzed using appropriate non-parametric statistical models (Kruskal-Wallis using Conover-Inman to make all pairwise comparisons between groups) with StatsDirect software v. 2.7.8. and compared to pretreatment and vehicle controls.

### Mouse lung infection model

*Ethical statement:* All animal experiments were performed under UK Home Office License 40/3644, and with local ethical committee clearance (The University of Manchester AWERB). All experiments were performed by technicians who had completed at least parts 1 to 3 of the Home Office Personal License course and held current personal licenses.

NOSO-502 was tested against *K. pneumoniae* NCTC 13442 (expresses OXA-48 carbapenemase, MIC= 1 μg/ml) in a neutropenic mouse pulmonary infection model by Evotec (Manchester, UK). Male CD-1/ICR mice 6-8 weeks old (Charles River UK) were allowed to acclimatize for 7 days, then rendered neutropenic by IP injection of cyclophosphamide (200 mg/kg on day 4 and 150 mg/kg on day 1 before infection). Mice were infected by intranasal route (∼4 × 10^6^ CFU/mouse) under parenteral anaesthesia. At 2 h, 8 h, 14 h and 20 h post infection, mice received treatments with NOSO-502 at 2, 6 or 20 mg/kg or with tigecycline at 40 mg/kg administered by SC route in a volume of 10 mL/kg (8 mice per dose). At 2 h post infection NOSO-502 was delivered once by SC route at 80 mg/kg in a volume of 10 mL/kg (8 mice). At 2 h post infection, one infected group was humanely euthanized, and lungs processed for pre-treatment quantitative culture to determine *Klebsiella* burdens. At 26 h post infection, all remaining mice were humanely euthanized. Lungs were aseptically removed, homogenized, serially diluted, and plated on CLED (cystine lactose electrolyte deficient) agar for CFU titers.

## ACKNOWLEDGMENTS

Some of the research leading to these results was conducted as part of the ND4BB ENABLE Consortium and has received support from the Innovative Medicines Initiative Joint Undertaking under Grant no 11583, resources of which are comprised of financial contributions from the European Union’s seventh framework program (FP7/2007-2013) and EFPIA companies’ in-kind contribution. The authors would like to thank Douglas Huseby, Diarmaid Hughes, Sha Cao, Richard Svensson, and Pawel Baranczewski from Uppsala University, Edgars Liepins, and Solveiga Grinberga from the Latvian Institute of organic synthesis for their contributions.

